# Towards a Taxonomy Machine – A Training Set of 5.6 Million Arthropod Images

**DOI:** 10.1101/2024.07.15.600863

**Authors:** D Steinke, S Ratnasingham, J Agda, H Ait Boutou, I Box, M Boyle, D Chan, C Feng, SC Lowe, JTA McKeown, J McLeod, A Sanchez, I Smith, S Walker, CY-Y Wei, PDN Hebert

**Affiliations:** Centre for Biodiversity Genomics, University of Guelph, 50 Stone Road East, Guelph, Ontario, N1G 2W1, Canada; Department of Integrative Biology, University of Guelph, 50 Stone Road East, Guelph, Ontario, N1G 2W1, Canada; Vector Institute, Toronto, Ontario, M5G 1M1, Canada

## Abstract

The taxonomic identification of organisms from images is an active research area within the machine learning community. Current algorithms are very effective for object recognition and discrimination, but they require extensive training datasets to generate reliable assignments. This study releases 5.6 million images with representatives from 10 arthropod classes and 26 insect orders. All images were taken using a Keyence VHX-7000 Digital Microscope system with an automatic stage to permit high-resolution (4K) microphotography. Providing phenotypic data for 324,000 species derived from 48 countries, this release represents, by far, the largest dataset of standardized arthropod images. As such, this dataset is well suited for testing the efficacy of machine learning algorithms for identifying specimens to higher taxonomic categories.

## Summary

The identification of organisms is a fundamental part of recognizing and describing biodiversity. The development of automated methods, which can identify specimens without involving taxonomists, is critical given the taxonomic impediment [1,2]. One effective solution involves the adoption of identification systems based on the analysis of sequence variation in short, standardized DNA regions [3]. As well, digitization initiatives at major natural history collections [6] and imaging linked to large DNA barcoding projects [4,5] are providing the basis for image-based identification systems driven by machine learning algorithms.

Images can be used to build classification systems capable of identifying species [7,8,9]. Various methods and datasets have been proposed to advance image-based identifications for arthropods [10,11]. Most past work has considered arthropod classification from: 1) the context of integrated pest management [12,13,14], 2) as part of crowd-sourced citizen science efforts such as iNaturalist [15,16] or 3) interpreting data acquired by camera traps [17,18].

Machine learning algorithms, especially convolutional artificial neural networks and their variants have emerged as the most effective method for object recognition and detection [10,19,20]. However, data sets of sufficient size to properly train these models are scarce [20,21,22,23]. In recent years, substantial efforts have been invested into the digitization of natural history collections which have imaged both individual specimens [24,25,26] and entire drawers of specimens [6,27,28]. This work provides training material for classification algorithms; but the images are structurally uniform. Specimens are usually mounted and therefore captured in a single standard orientation (mostly dorsal view) limiting their application in less regimented contexts [29]. In addition, there are biases in selecting insects which translate into the availability of specimens for digitization or images gathered by community science [30,31].

In this study, we present a dataset of 5.6 million arthropod images gathered during a large-scale DNA barcoding project. As specimens were not placed in a standard orientation before photography, any machine learning-based object detection must interpret specimens in varying orientations to classify them taxonomically, an approach which requires massive training sets.

## Data generation

The present dataset was generated as part of an ongoing effort to build a global DNA barcode reference library for terrestrial arthropods [32]. The workflow involves placing each small (<5 mm) specimen into a well of a round-bottom 96-well microplate before DNA extraction. Specimens are individually photographed at 4K resolution in plate format using a Keyence VHX-7000 Digital Microscope system with a fully integrated head and an automatic stage (Figure 1). The setup uses an adjustable Keyence illumination adapter to ensure uniform light conditions and a scanning stage equipped with a custom-engineered mount that holds each plate (Figure 1, Supplementary File 1). This system can capture 95 images within 15 minutes by controlling stage movements in X-Y coordinates. It also has the capability to automatically control the height of the stage with a precision of 0.1 μm. By moving the lens throughout the different focal planes of each specimen, the VHX system captures every pixel that comes into focus at each level and combines them into a single, fully focused image. For all images the system was set to a brightness of 27.8ms and colours to R1.7 G1.0 B2.19.

**Figure 1:**
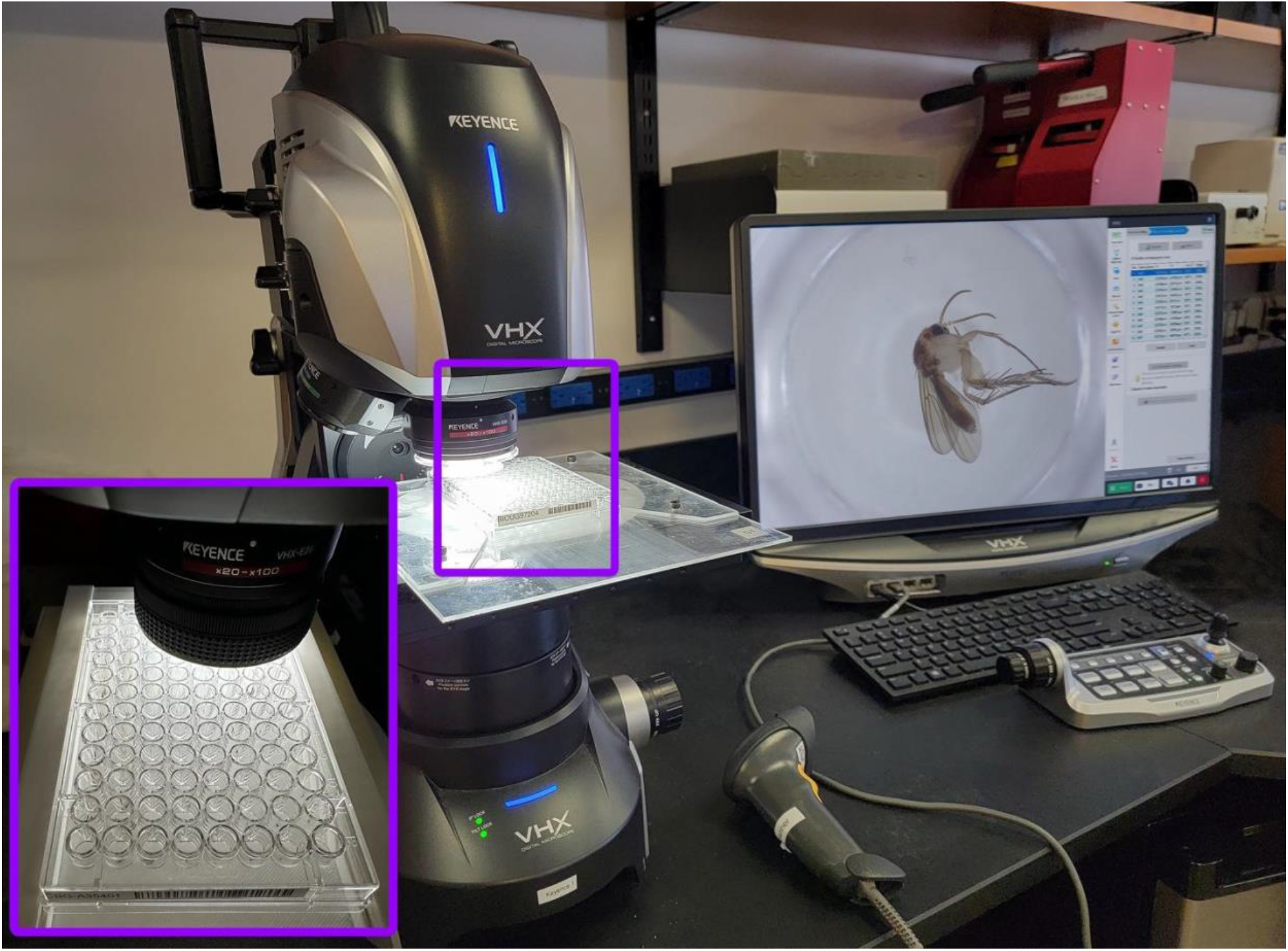
Keyence VHX-7000 Digital Microscope system. The inset shows a microplate within the custom-engineered mount.

The resulting images are packaged with a Python script (Supplementary File 2) and uploaded to the Barcode of Life Datasystems (BOLD) [33] where they are automatically associated with individual specimen records. A backup copy of each image is subsequently transferred to a third-party cloud service using an automated script (Supplementary File 3).

## Data Description

The present dataset includes 5,675,731 images of mostly terrestrial arthropods. It includes 1.13 million images from a previous release [20] together with 4.54M images generated from 2019-2022. The image size is 2880 × 2160 pixels, creating an average file size of 17.9MB for a tif and 1.88 MB for a jpg. Figure 2 shows an array of 95 images taken with the Keyence system (the empty space at the lower right corresponds to an empty control well in the microplate).

**Figure 2:**
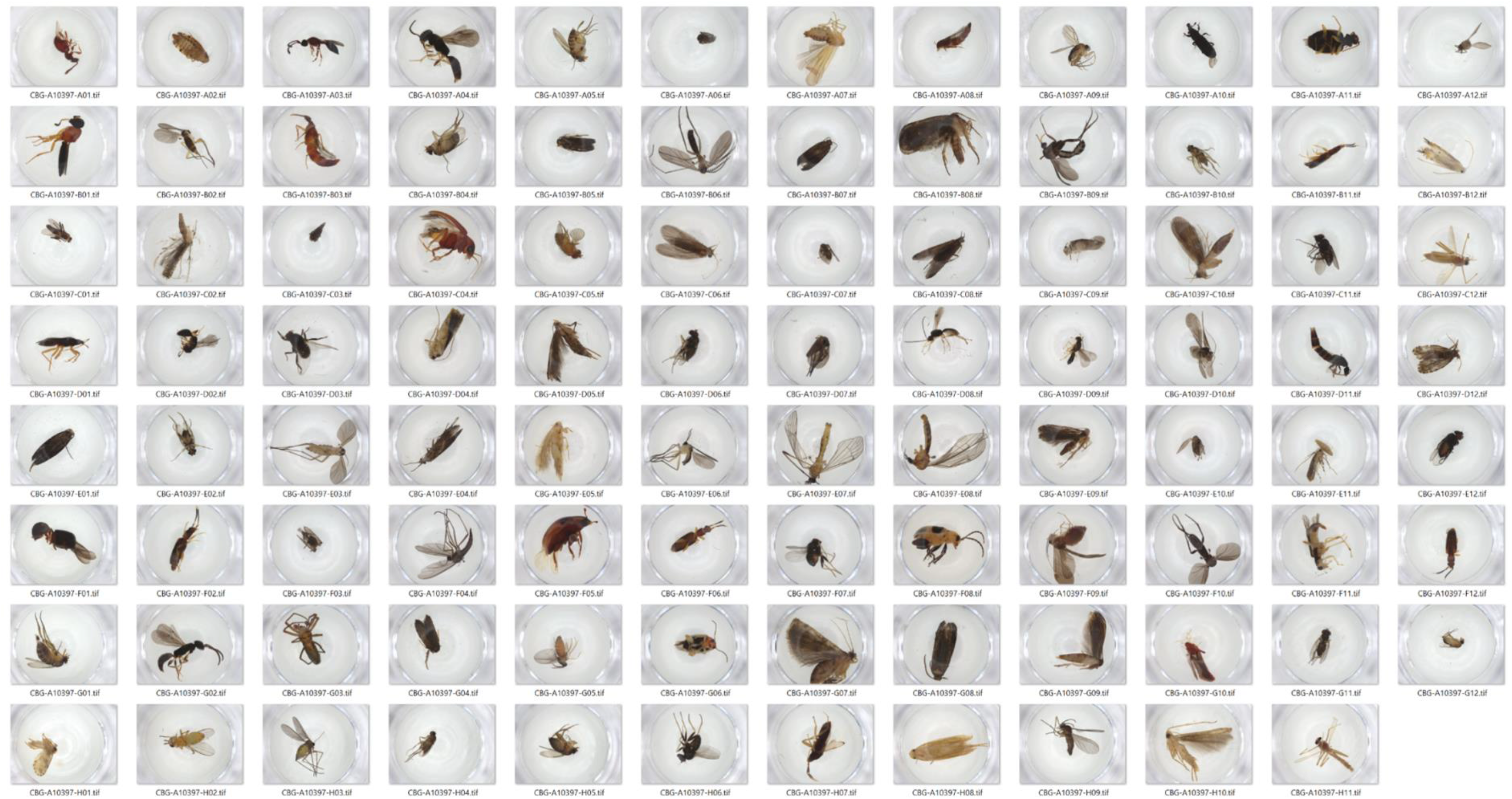
Panel of example images taken with the Keyence setup. The empty space in the lower right corresponds to an empty control well in the microplate.

### Geographic coverage

The dataset contains images for specimens from 1,698 sites in 48 countries (Figure 3A). Most specimens derived from Costa Rica (62%), followed by South Africa (6%), United States (5%), and Thailand (3%) (Figure 3B, Supplementary File 4). Most specimens were collected using Malaise traps, but about 1,500 were captured using plankton and dip nets [34].

**Figure 3:**
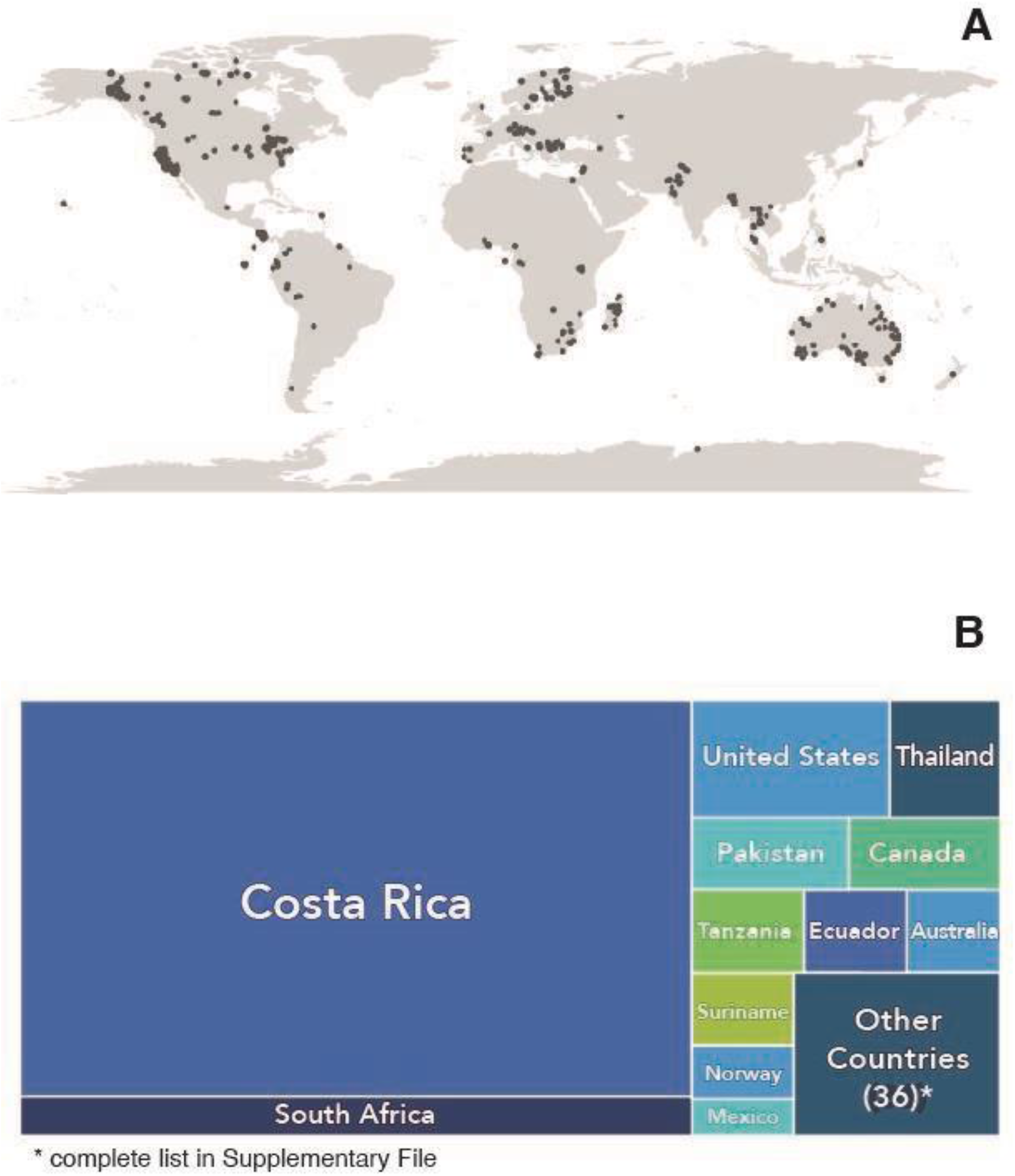
A) Sampling locations for photographed specimens. B) Tree map of countries of origin.

### Taxonomic coverage

Most images show individuals from the class Insecta (98%) (Figure 4A). Figure 4B shows the distribution of insect orders within the dataset. A very high proportion of all specimens have taxonomic assignments at the ordinal (99.4%) and family (85.9%) levels, but only 20.3% possess a generic assignment. Just 7.6% of the specimens (430,036) have a Linnean species designation, but 90.6% (N=5,143,970) possess a BIN assignment (Barcode Index Number, a species proxy) [35]. A total of 324,427 BINs are represented in this dataset.

**Figure 4:**
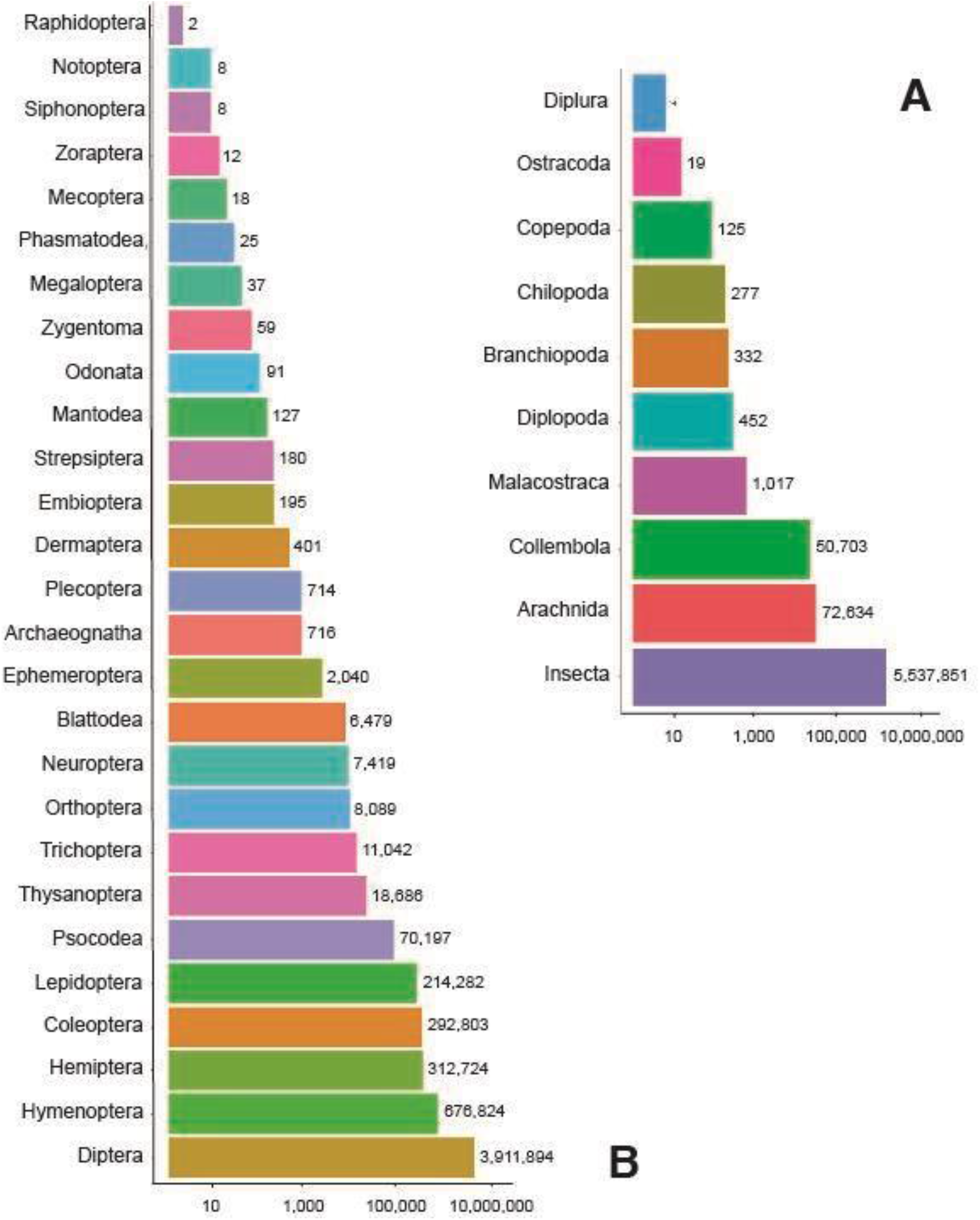
Log-scale plot showing coverage for 10 arthropod classes (A) and 27 insect orders (B).

## Conclusions

This dataset of 5.6 million images representing over 320,000 species of arthropods, mostly insects, is the largest of its kind. It is also unique in the geographic distribution of the photographed samples. As such, it represents a dataset which should aid the development of machine learning algorithms for object recognition and classification. These images can also be used to estimate biomass [36,37] or even abundance when entire communities are digitized [38,39]. Each image in the dataset is also available as an element of the collateral data for a barcode record on BOLD [33] providing support for taxonomic assignment and to enable direct visual comparisons between individuals.

By using three Keyence VHX-7000 systems for 50h per week, the Centre for Biodiversity Genomics generates three million images per year. The deployment of Keyence systems at a few core facilities could readily generate ten million images per year, allowing rapid growth in the training sets required to hone AI-enabled identification systems.

## Acknowledgements

All images were generated at the Centre for Biodiversity Genomics as one component of an integrated program to construct a DNA barcode reference library. As such, the assembly of this dataset benefitted from the contributions of all CBG staff as well as colleagues around the world who are aiding the advance of this work. We thank Suzanne Bateson for graphics support.

## Data availability statement

The dataset and a separate metadata file in *.tsv format (Supplementary File 4 contains definitions of all metadata fields) can be accessed at https://biodiversitygenomics.net/5M-insects/.

